# Biocompatible Multi-functional Polymeric Material for Mineralized Tissue Adhesion

**DOI:** 10.1101/2025.05.30.656989

**Authors:** Yan Luo, Chenyang Zhang, Sage Fulco, Jingyi Liu, Keyu Chen, Yuntao Hu, Yuchen Jiang, Rui Xu, Leela Rakesh, Ozer Fusun, Ottman Tertuliano, Kevin Turner, Kyle H. Vining

## Abstract

This study developed a biocompatible multifunctional thiol-ene resin system for adhesion to dentin mineralized tissue. Adhesive resins maintain the strength and longevity of dental composite restorations through chemophysical bonding to exposed dentin surfaces after cavity preparations. Dental pulp cells are exposed to residual monomers transported through dentinal tubules. Monomers of conventional adhesive systems may result in inhomogeneous polymer networks and the release of residual monomers that cause cytotoxicity. In this study, we develop a one-step multi-functional polymeric resin system by incorporating trimethylolpropane triacrylate (TMPTA) and bis[2-(methacryloyloxy)ethyl] phosphate (BMEP) to enhance both mechanical properties and adhesion to dentin. Molecular dynamics simulations identified an optimal triacylate:trithiol ratio of 2.5:1, which was consistent with rheological and mechanical tests that yielded a storage modulus of ~30 MPa with or without BMEP. Shear bond tests demonstrated that the addition of BMEP significantly improved dentin adhesion, achieving a shear bond strength of 10.8 MPa, comparable to the commercial primer Clearfil SE Bond. Nanoindentation modulus mapping characterized the hybrid layer and mechanical gradient of the adhesive resin system. Further, the triacrylate-BMEP resin showed biocompatibility with fibroblasts in vitro. These findings suggest the triacrylate-trithiol crosslinking and chemophysical bonding of BMEP provide enhanced bond strength and biocompatibility for dental applications.

## Introduction

Clinical applications involving bone and dental tissues require polymeric adhesives for etching and bonding to mineralized tissues. Conventional bone fixation methods, such as metal implants for bone fractures, show limitations in complex cases involving small, unstable fragments^1^. In restorative dentistry, failed restorations and tooth fractures remain a significant problem despite advancements in dentin resin adhesive materials^2, 3^. Further, residual monomers released from adhesive materials can negatively impact surrounding biological tissues.^4, 5^ There remains a need for biocompatible polymeric materials that can achieve strong adhesion to mineralized tissues.

Dental decay is increasing in prevalence, with over 3 billion untreated cases, despite increased access to prevention and dental treatments worldwide.^6^ Dental decay is treated by removing the decayed tooth material, preparing the tooth for direct access to dentin, and placing a direct resin restoration with a bonded adhesive resin interface.^7^ Adhesion to the dentin protects the restoration from microleakage, tooth fracture, infection, and ultimate failure of the treatment.^8^ The widely used commercial methacrylate-based adhesive, Clearfil SE Bond, only shows 56% double bond conversion for the methacrylate functional groups^9^. Lack of conversion may lead to inhomogeneous polymer networks, residual monomers in the resin, and cytotoxicity and disruption of cellular signaling ^4, 10^. Hydrolytic degradation and water sorption compromise the long-term durability of adhesive resins ^10–12^. Acrylamide-based polymers have been developed to improve bond stability and resistance to enzymatic and hydrolytic degradation.^13, 14^

Thiol-ene click chemistry has been investigated for bonding to mineralized tissues with rapid curing, minimal residual monomer content, and high cytocompatibility^15^. Thiol-ene-based polymeric materials can achieve mechanical strength comparable to methacrylate and polyurethane systems while minimizing cytotoxic effects.^15^ Trimethylolpropane triacrylate (TMPTA) and structurally-related tri-thiol (TMPTMP) monomers are investigated as three-armed thiol-ene crosslinked polymers for a biocompatible resin.^16^ Three acrylate groups in TMPTA provide a high degree of crosslinking, which improves thermal stability, mechanical strength, and glass transition temperature in UV-curable systems^17, 18^. TMPTA also enhances rheological properties like shear viscosity and storage modulus in various polymer applications, offering fast curing times and improved material hardness, making it more efficient than other monomers^19, 20^. Thiol-ene materials crosslinked with TMPTA supported the adhesion, proliferation, and differentiation of dental pulp stem cells (DPSCs), providing a bioinstructive environment for tissue regeneration.^16^

Bis[2-(methacryloyloxy) ethyl] phosphate (BMEP) is investigated in this study as an adhesive monomer. The phosphate group in BMEP reacts with mineralized tissues, such as bone and dentin, by interacting strongly with hydroxyapatite (HA), enabling effective demineralization, the formation of longer resin tags, and a stable hybrid layer ^2, 21, 22^. This interaction promotes deeper etching compared to other adhesive systems like 10-MDP, which primarily rely on superficial ionic bonding^10, 12, 22^ and enable an acid-free etching procedure. The methacrylate groups of BMEP can react with thiol-ene click crosslinking, promoting a high degree of polymerization to strengthen the adhesive network and enhance its mechanical properties^15, 23^. Furthermore, BMEP acts as an effective matrix metalloproteinase (MMP) inhibitor, protecting the collagen matrix from enzymatic degradation and improving the long-term durability of the resin-dentine bond^11, 23, 24^. Additionally, compared to other etching systems, BMEP’s ability to perform both chemical and micromechanical retention makes it superior in creating a robust adhesive interface^12, 25^. This study aimed to develop a biocompatible adhesive with strong dentin adhesion based on the multifunctional properties of TMPTA and BMEP.

## Experimental Methods

### Materials

Chemicals used in this study, Trimethylolpropane triacrylate (TMPTA), Trimethylolpropane tris(3-mercaptopropionate) (TMPMP), Bis[2-(methacryloyloxy)ethyl] phosphate (BMEP) and 2,2-Dimethoxy-2-phenylacetophenone (DMPA) were all purchased from Sigma-Aldrich. All chemicals in experiments were used as received without further purification.

### Material Fabrication

The polymeric resin material was prepared using a dual monomer system consisting of TMPTA and TMPMP (**Figure 1a**), utilizing thiol-ene click chemistry (**Figure S1**), which facilitates rapid polymerization through a radical-mediated mechanism, and the photoinitiator, DMPA, was incorporated into the resin formulation to initiate the light-curable reaction. A range molar ratios of TMPTA and TMPMP was tested (1.5:1 to 5.5:1). 5% w/v BMEP was used for samples containing BMEP and 0.05% w/v DMPA was immediately added prior to shaker mixing for several minutes for homogenization. The solution was mixed at room temperature in a clean amber vial to prevent direct ambient light exposure.

**Figure 1.**
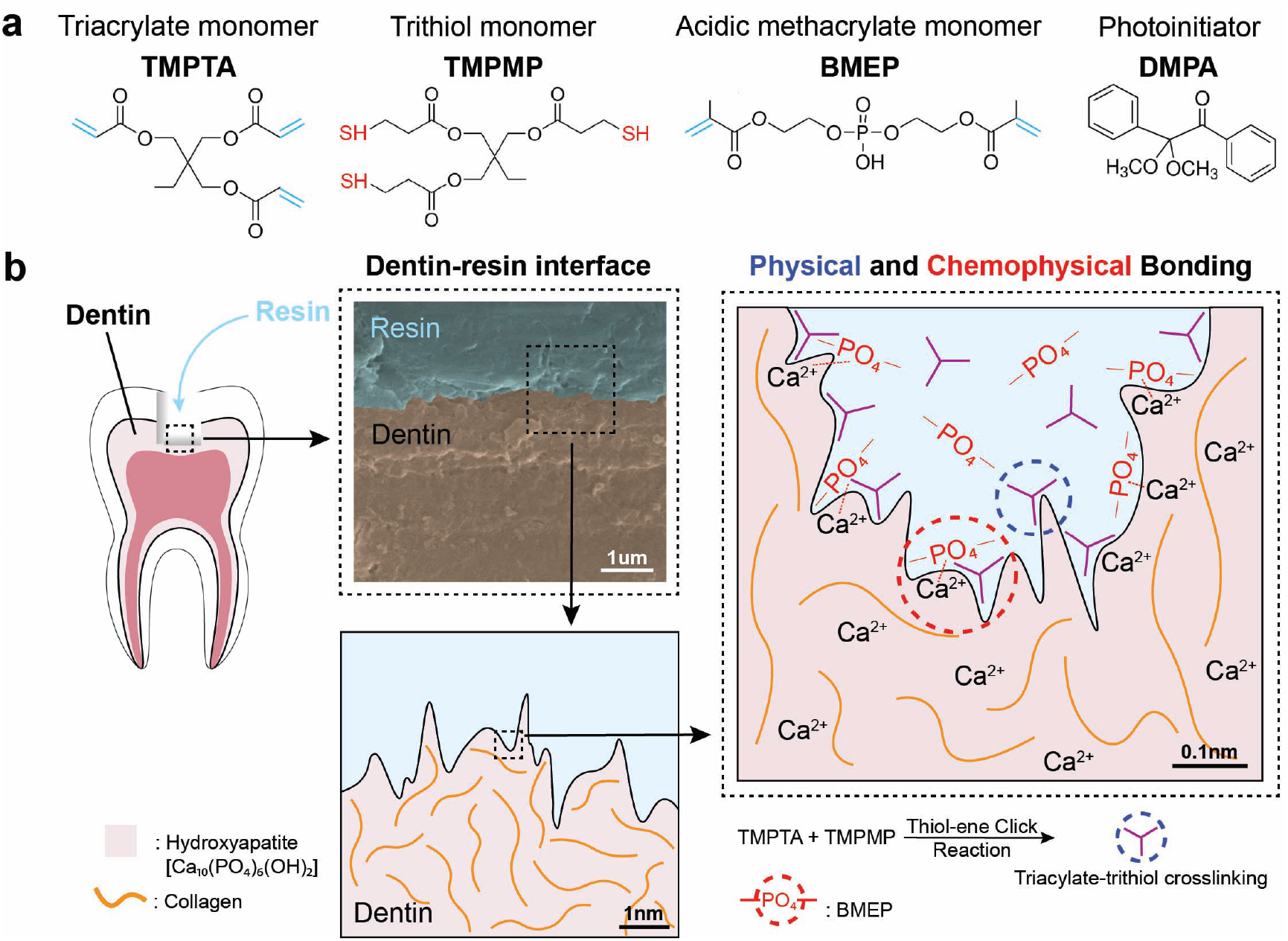
Illustration of TMPTA-TMPMP resin-dentin dual adhesion mechanisms. **(a)** Overall schematic of resin with Trimethylolpropane triacrylate (TMPTA) and Trimethylolpropane tris(3-mercaptopropionate) (TMPMP) as monomers, Bis[2-(methacryloyloxy)ethyl] phosphate (BMEP) as the primer, and 2,2-Dimethoxy-2-phenylacetophenone (DMPA) as the photoinitiator. **(b)** Demonstration of both physical and chemophysical interlocks between resin and dentin through crosslinked polymers and chelation reactions involving BMEP.

### Molecular Dynamics Simulation

Molecular dynamics (MD) simulation was conducted using Blend in Material Studio (http://accelrys.com/products/collaborative-science/biovia-materials-studio/). The process entails constructing desired chemical molecules, optimizing their minimum energy utilizing a preferred force field, and employing the Blend module for compatibility assessments. For example, it effectively evaluates the compatibility of the desired chemical molecule TMPTA with TMPMP. Throughout calculations, critical parameters such as temperature dependence of interaction (*χ*), binding energy, and phase diagrams are meticulously computed using theories like Gibbs free energy and Flory-Huggins. The outcome is a comprehensive table generated through energy minimization techniques, identifying the lowest energy pairs. Additionally, overlays assist in pinpointing the lowest energy absorption sites, revealing numerous compelling results. Once all molecules are individually minimized, blend computations are initiated by selecting base molecules (TMPTA) and screen molecules (TMPMP). The binding and potential energy data were analyzed of different combinations of TMPTA and TMPMP with simulations up to 10,000 steps during the production run. These calculations are executed at room temperature and atmospheric pressure, determining interaction energies for different mixtures. The chosen blending method provides crucial information on binding energy, coordination numbers, Flory-Huggins interaction parameter (*χ*), mixing, and interaction energy.

### Oscillatory Shear Rheology

Rheology data were obtained using an HR30 Discovery Hybrid Rheometer (TA Inc.). 80 μL of the resin solution was dispensed onto the loading plate with the trim gap set to 1000 nm. Then, a time sweep of oscillatory shear rheology with strain % of 10^−3^ and a frequency of 1.0 Hz was carried out at 25°C for a duration of 930 seconds (30 seconds delay followed by 900 seconds of data collection). The power density of the ultraviolet (UV) source (Omnicure S2000, Excelitas) was 0.25 mW/cm^2^. The geometry gap was set to float to maintain a constant normal force. Polymerization shrinkage is measured by the change in gap during curing, defined by 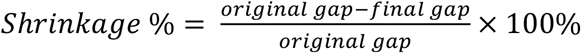.

### Scanning Electron Microscopy (SEM)

Dentin samples were prepared by removing enamel from human molars using IsoMet Low Speed Saw, and the cutting edge was sanded with 150 and 220 grit sandpapers. The resultant dentin samples had cuboid shapes, and resin solution was pipetted onto one of the cuboid surfaces which had the largest surface area. A cover glass treated with Smooth-On Ease release spray was positioned over resin solution to avoid oxygen inhibition. After being cured for 60 seconds under UV light with an intensity of 103.2 mW/cm^2^, the dentin sample coated with resin was detached from the cover glass. The interface between dentin and resin was further sanded with 150 and 220 grit sandpapers. Subsequently, SEM imaging of dentin samples was performed using Quanta 600 FEG ESEM.

### Shear Bond Strength Test

Human third molars were obtained by from the Penn Dental Oral Surgery clinic, following approve IRB protocol #827807. Dentin surfaces were prepared using a 600-grit polishing wheel (Buehler Ltd., Lake Bluff, IL, USA) and subsequently embedded in acrylic resin blocks. Resin solutions were applied to the denting surfaces in cylindrical plastic molds with the contact area being 2.1 mm^2^ and cured. All shear tests were carried out with a universal shear bond tester (BISCO Inc., Schaumburg, IL, USA) applying a force (N) parallel to the interface between the dentin and adhesive. Shear bond strength was calculated in megapascals using the formula: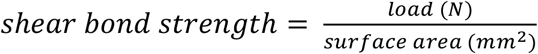.

### Nanoindentation Mapping

Nanoindentation was performed on a TI-950 nanoindenter (Hysitron, USA) with a diamond Berkovich pyramidal tip. Hardness and modulus were measured as described by Oliver and Pharr ^26^, under load control, to a peak load of 8 mN, with a 5-second loading time, 2-second hold time at the peak load, and 5-second unload time. A 25-nm liftoff was used before testing to allow for proper surface detection^27^, and the reduced elastic modulus was calculated from the unloading curve^26^. Testing was performed on the dentin-resin interfaces, with the indentation direction parallel to the interfacial plane. A testing array was constructed to measure the elastic modulus of the dentin and resin as a function of the distance from the interface.

### Biocompatibility Test

Resin samples with and without BMEP were soaked in low glucose DMEM (Gibco) at 37°C with 5% CO_2_ for 7 days. Collected condition media were diluted with low glucose DMEM to a final concentration of 100%, 50%, and 25% of the original condition media. BJ cells were seeded at 62500 cells cm^-2^ in 96 well-plate and incubated in low glucose DMEM (Gibco) with 10% fetal bovine serum (Cytiva). After 24 h, cell culture media were replaced by 100 *μ*L serially diluted original condition media, and cells were kept culturing for 24 h. For fluorescence imaging, cells were fixed in 2% PFA for 30 min and then stained with Phalloidin and DAPI. For cell viability experiments, released LDH in cell culture supernatants was measured by CytoTox 96® Non-Radioactive Cytotoxicity Assay (Promega). Relative cell viability was compared with maximum LDH release control. For cell counting experiments, cells were trypsinized by Trypsin-EDTA (0.05%) (Gibco) and retrieved from 96 well-plate. Retrieved cells were stained with 0.4% solution of trypan blue for counting.

### Statistical Methods

GraphPad Prism was used for statistical analysis and data visualization. Rheology data was analyzed with unpaired t test. Shear bond test data was analyzed with Brown-Forsythe and Welch ANOVA test. Sample size and p-values are noted in figures. Cell viability and cell counting data were analyzed with 2-way ANOVA test.

## Results

The multi-functional resin material features physical and chemophysical adhesion mechanisms for bonding with mineralized tissues like dentin (**Figure 1)**. The roughness and unevenness of the dentin surface provide micromechanical, physical bonding between the triacylate-trithiol network and micro-roughness on the interface. Monomer BMEP provides chemophysical bonding by chelation of calcium ions in hydroxyapatite of dentin by the phosphate group in BMEP. BMEP can also etch the surface, which increases roughness for physical bonding. Therefore, BMEP enables a one-step resin adhesive system.

Cross-linked polymer networks reinforce the physical bonding of resin to dentin and increase the cohesive strength of the material (**Figure 2a**). The cross-linking of TMPTA polymers was investigated by molecular dynamics (MD) simulations and oscillatory rheology. MD calculations of TMPTA and TMPMP simulated the resin materials at the molecular level to predict polymerization dynamics and crosslinking efficiency^28^. The red solid curves show the energy between base materials and screening materials (U_BS_), the blue solid curves are the energy between base materials and base materials (U_BB_), and the black dotted curves represent the energy between screening materials and screening materials (U_SS_) (**Figure 2b**). The results were evaluated to identify the appropriate reacting ratios by 1) minimizing U_BS_, 2) aligning interaction energies, and 3) finding the peak of U_BS_ in between of U_SS_ and U_BB_. U_BS_’s potential energy is relatively low with molar ratios of 1.5:1 to 3:1. Lower potential energy of U_BS_ in atomistic simulations should indicate the mechanical stability of the resin ^28^. The red dashed lines indicate the lowest U_BS_ peak potential energy across six ratios. With increasing TMPTA and TMPMP molar ratios, the axis-binding energy difference between U_BS_ and U_SS_ decreased while the difference between U_BS_ and U_BB_ increased (**Figure 2c**). Ratios ranging from 0.5:1 to 2.5:1 show both values near zero. U_SS_, U_BB_ and U_BS_ should have similar curve shapes, and their axis should approximately align with each other (namely, their axis’s position should have the same axis-binding energy value on x-axis). Aligning the interaction energies of different monomers in thiol-ene systems optimizes polymerization dynamics by facilitating molecular collisions along favorable reaction pathways^29^. The potential energy difference between U_BS_ and U_SS_ decreased and the difference between U_BS_ and U_BB_ increased (**Figure 2d**). The energy difference was largest with ratios at 0.5:1 and from 2.5:1 to 3:1. The peak of U_BS_ should be in between U_SS_ and U_BB_ and the one with larger energy differences is preferred because energy differentials between interacting components are critical for optimizing crosslinking reactions as larger energy differences drive more efficient polymerization^15^. Together, these results suggest that TMPTA: TMPMP ratio at 2.5:1 is most favorable for the thiol-ene reaction. The predicted molecular structure is shown in **Figure S2**.

**Figure 2.**
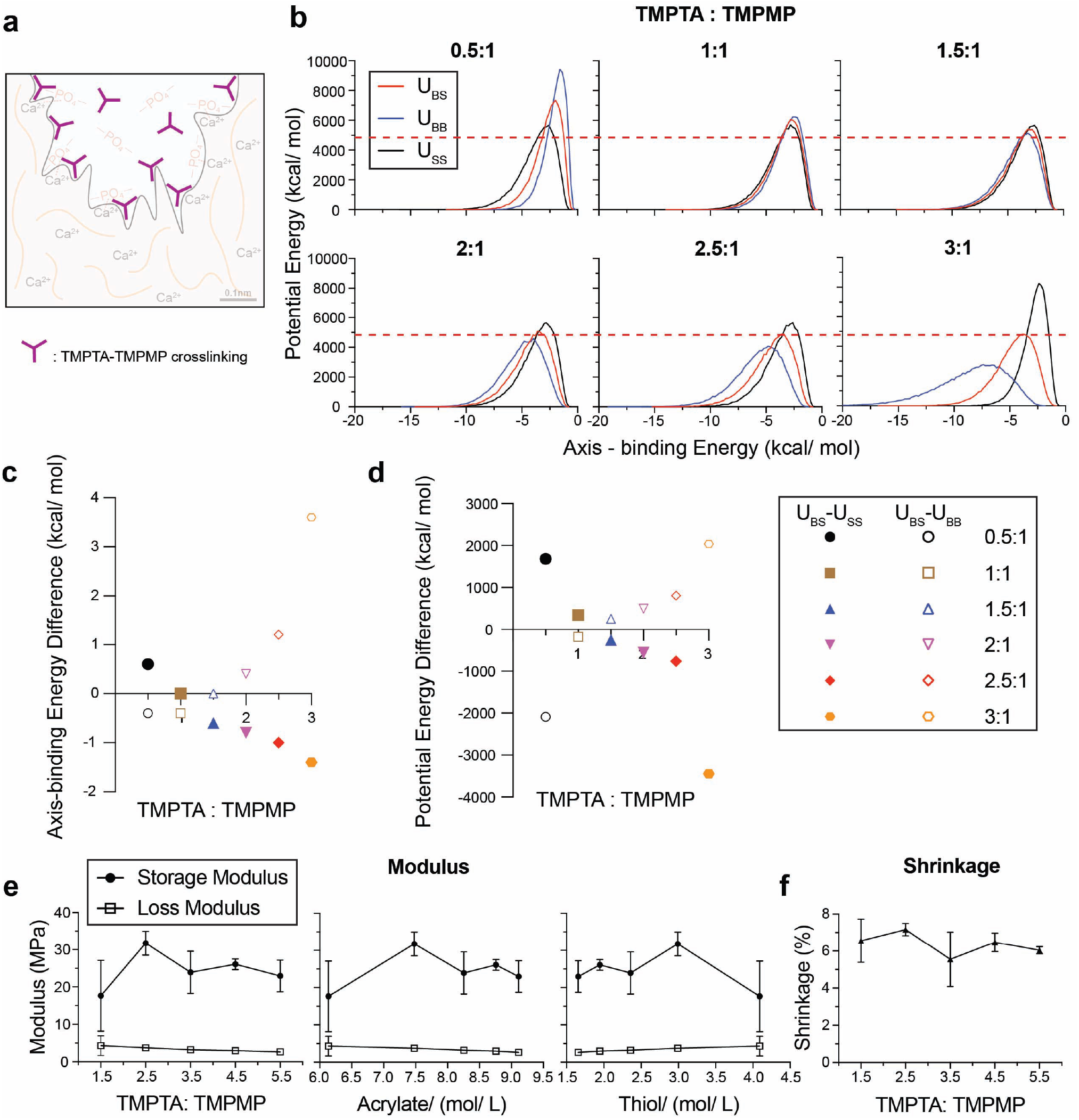
Molecular dynamics simulations and comparative modulus and shrinkage analysis for resin systems with varying compositions of TMPTA and TMPMP. **(a)** Illustration of TMPTA-TMPMP crosslinking in resin. **(b)** Molecular dynamics simulation results of potential energy and axis-binding energy profile for TMPTA and TMPMP reacting under different ratios. U – potential energy; S – screening material, TMPMP; B – base material, TMPTA. Summary plots for **(c)** axis-binding energy difference and **(d)** potential energy difference between UBS and USS or UBB for different TMPTA: TMPMP ratios. **(e)** Storage modulus and loss modulus of resin plotted as a function of: TMPTA: TMPMP ratios, acrylate concentration and thiol concentration. **(f)** Shrinkage of resin under different TPMTA: TMPMP ratios.

Oscillatory shear rheology was performed on resins with TMPTA: TMPMP ratios ranging from 1.5: 1 to 5.5: 1 with or without BMEP. Storage and loss moduli plotted with acrylate and thiol concentrations showed the highest storage modulus at 7.5 mol/L TMPTA concentration and ~2.9-3.0 mol/L TMPMP concentration, corresponding to a 2.5 molar ratio of TMPTA: TMPMP (**Figure 2e**). And shrinkage remained stable at around 6% through all ratios (**Figure 2f**). These data showed that increasing acrylate concentration above 7.5 mol/L (ratio=2.5) resulted in a trend of decreasing storage modulus, which is consistent with the MD simulations in **Figure 2b-d**.

The storage moduli of resin samples with BMEP (5 wt%) ranged from 23 to 36 MPa and remained slightly higher across all ratios than samples without BMEP (17~32 MPa). BMEP did not have a significant impact on the loss modulus (~3.5MPa) and polymerization shrinkage (~6%) (**Figure S3**). These values are comparable to other resin systems based on thiol-ene click chemistry^15, 30, 31^ and polymerization shrinkage studies (1~6%)^32–34^.

We investigated the mechanical and adhesive properties of the triacrylate/trithiol resin (TMPTA: TMPMP = 2.5) at the dentin-resin interface. Nanoindentation was performed on a thin film of TMPTA/TMPMP resin to evaluate the homogeneous curing and nanomechanical properties. The thiol-ene TMPTA and TMPMP radical-initiated propagation rate constant was estimated to be nearly 1000 times higher than that of methacrylate-methacrylate groups^28^, which leads to a rigid and homogeneous covalently crosslinked polymer network. This homogeneity is verified through the nanoindentation modulus in **Figure S4**. It shows the modulus was uniformly around 1GPa across the resin film’s frontside (0.992 GPa) and backside (1.125 GPa), with a higher modulus on the backside due to less oxygen inhibition^35^.

The adhesive interface between the human dentin and the resin material was characterized by SEM, EDS, shear bond strength, and shear rheology (**Figure 3**). The resin was cured onto dentin surfaces with exposed dentinal tubules. SEM imaging (**Figure S5**) showed resin tags in dentinal tubules, which was confirmed by the sulfur elemental analysis (yellow) (**Figure 3b**). When cured onto smooth, polished dentin surfaces, SEM imaging on the resin layer (196 *μ*m thick) showed a smooth interface with dentin with no defects, suggesting an intact micromechanical bonded interface (**Figure 3c**). Shear bond tests were performed on smooth, polished dentin substrates (**Figure 3d**). The average shear strength of resin materials with and without BMEP was 10.8 MPa and 1.5 MPa, respectively (**Figure 3e**), compared to 13.3 MPa with the commercial primer Clearfil SE Bond. The adhesive strength of the multifunctional resin with BMEP showed no statistically significant difference with Clearfil SE Bond, which was consistent with the reported shear bond strength of other commercial adhesive resin systems^36^. Further, the modulus and shrinkage comparison of resin materials with or without BMEP did not show a statistically significant difference at a molar ratio of TMPTA: TMPMP = 2.5 (**Figure 3f**). These data suggest that the addition of BMEP significantly strengthens the adhesion on the resin-dentin interface without negatively impacting the resin system’s mechanical properties.

**Figure 3.**
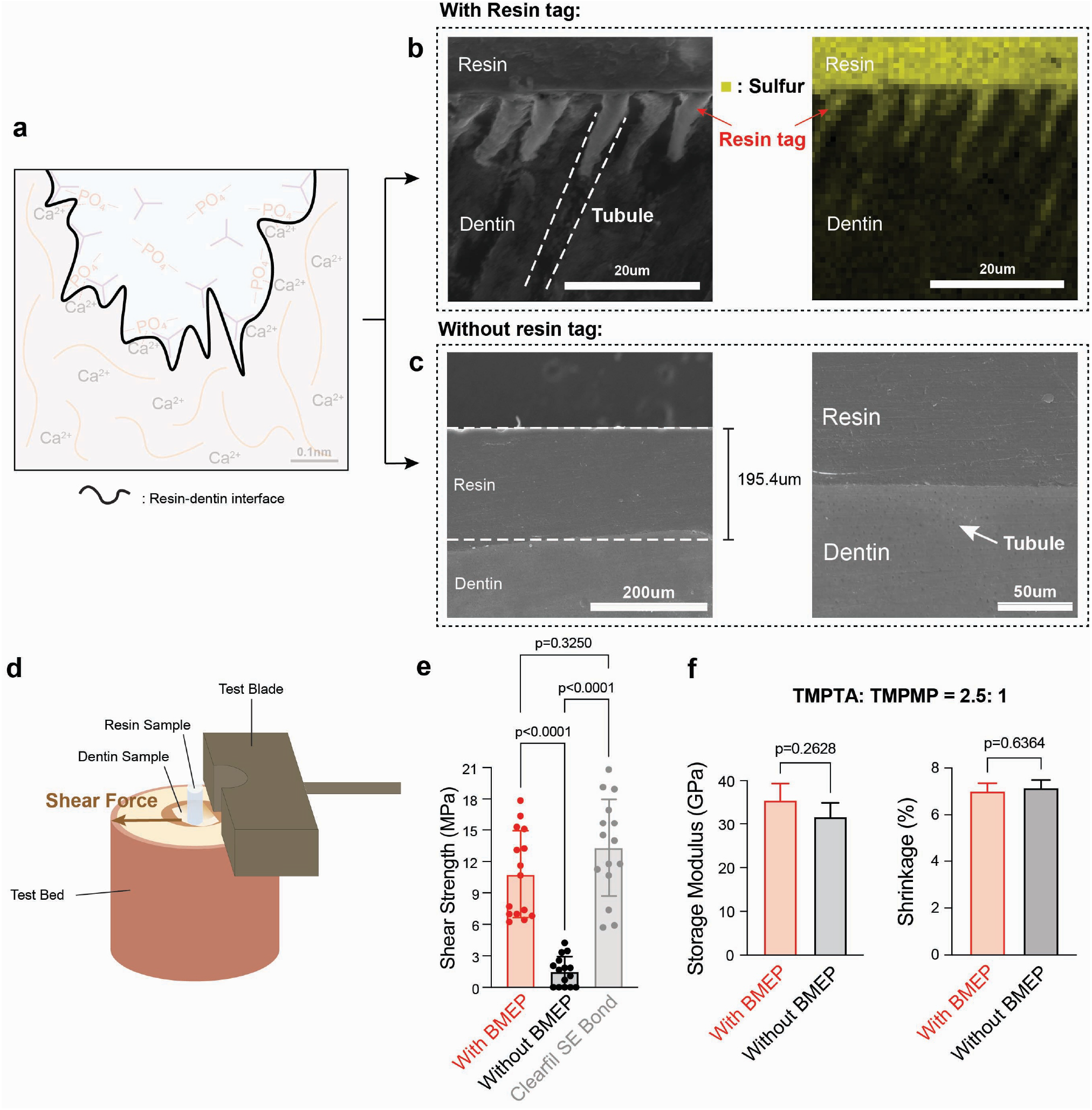
Comparative adhesion, storage modulus and shrinkage analysis of the resin-dentin interface and resin systems with and without BMEP. **(a)** Illustration of resin-dentin interface. **(b)** SEM (left) and EDS (right) images of dentin interface with resin tag exposed. (Yellow: sulfur). **(c)** SEM images of dentin interface without resin tag under 200 *μ*m and 50 *μ*m scale. **(d)** Shear test demonstration for adhesion situation between resin and dentin sample and **(e)** corresponding shear strength statistical results for resins and the commercial primer Clearfil SE Bond (Brown-Forsythe and Welch Test, sample size n = 15). **(f)** Statistical results of two resin systems about storage modulus and shrinkage percentage (unpaired t-test, sample size n = 3).

We hypothesized that the phosphate-hydroxyapatite chemophysical interactions of BMEP reinforce the nano-mechanical interface of resin and dentin (**Figure 4**). Nanoindentation line mapping was performed to investigate the nanomechanical properties of the dentin-resin interface and its hybrid layer with a line of 20 indentation tests (black dots), spaced 10*μ*m apart, beginning in the dentin 100*μ*m away from the interface and running to 100 *μ*m inside the resin (**Figure 4b**). Indents were spaced by a distance of 10 *μ*m to avoid any interference from subsurface plastic deformation from neighboring indents^27^. Improved spatial resolution across the interface of 1 *μ*m was achieved by running 10 separate lines of indents, each beginning 1*μ*m closer to the interface, with each line being spaced 10 *μ*m apart. This resulted in a total array of 200 indents on each specimen and an indentation test every 1 *μ*m away from the interface from 100 *μ*m inside the dentin to 100 *μ*m inside the resin.

**Figure 4.**
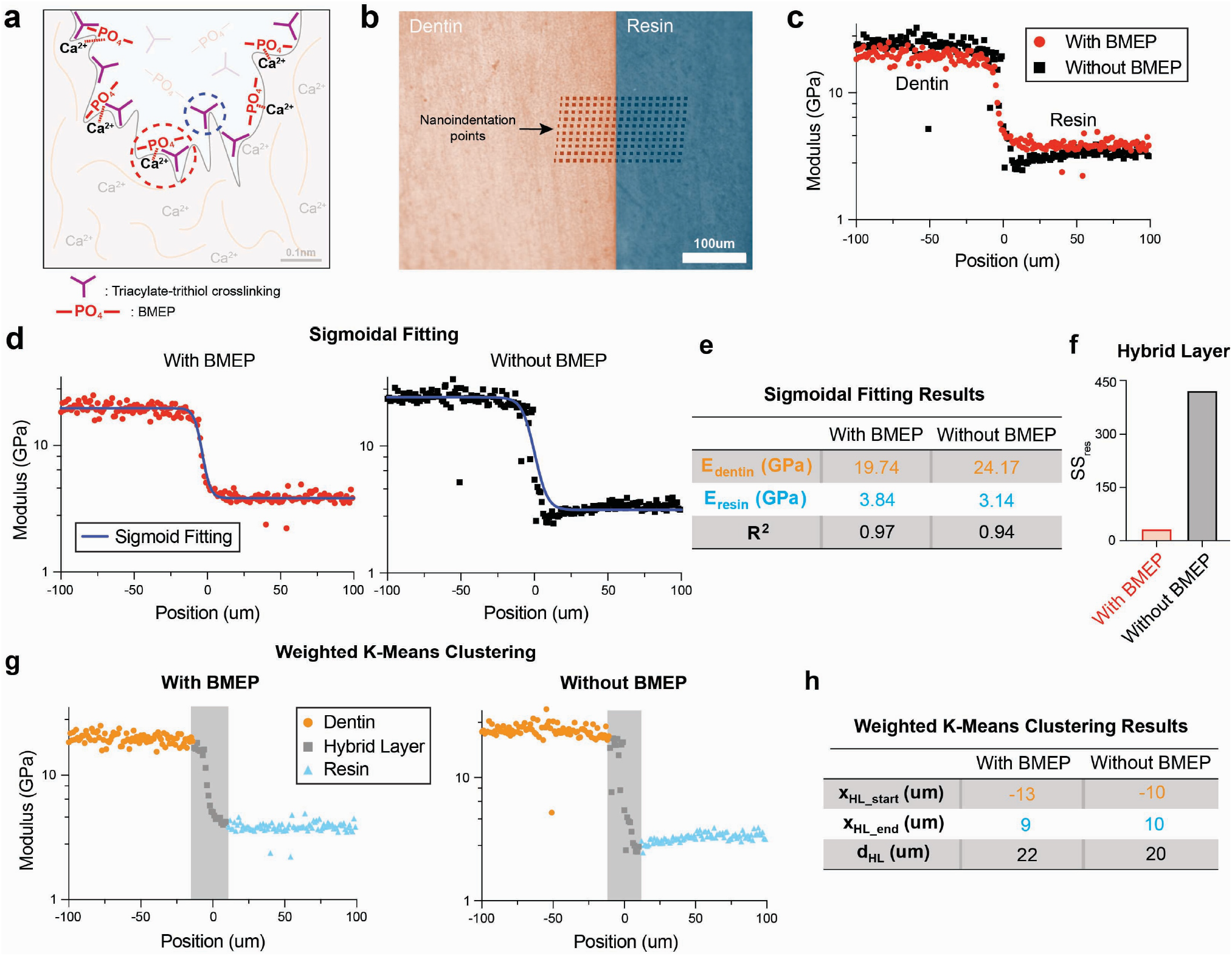
Nanomechanical analysis of the resin-dentin adhesive interface. **(a)** Illustration of two interlocking mechanisms on the resin-dentin interface. **(b)** Nanoindentation mapping array (10 sperate indentation lines * 20 indentation tests), **(c)** modulus results for resin with (red) and without (black) BMEP, plots for **(d)** sigmoidal fitting results (blue curves), **(e)** corresponding summary table, and **(f)** comparison on HL’s sum of residual (SSres). **(g)** Weighted k-means clustering results (dentin region – orange, hybrid layer (HL) region – grey, resin region – blue), and **(h)** corresponding summary table. All the results demonstrated here represent the aggregate outcomes derived from the analysis of all 10 individual indentation line tests.

Line mapping nanoindentation results of TMPTA/TMPMP resin samples with BMEP (red dots) and without BMEP (black squares) are shown in **Figure 4c**. Sigmoidal fitting was utilized to model the transition between dentin and resin modulus (**Figure 4d**) because a similar sigmoidal shape was observed in nanoindentation studies of the dentin-enamel junction ^37, 38^. The sigmoidal expression to describe the relationship between modulus and indentation position is 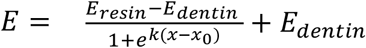. E represents modulus, k is a fitting parameter, x is the position, and x_0_ is the shifted origin. The fitting parameter results in **Figure 4e** show a 22% higher modulus (3.84 GPa, R^2^ = 0.97) with BMEP compared to without BMEP (3.14 GPa, R^2^ = 0.94). Further, **Figure 4f** shows that BMEP improved the sigmoidal transition of modulus from dentin to resin, quantified by a lower sum of squared residuals (SS_res_) within the hybrid layer (32.6), compared to resin without BMEP (422.4). Weighted k-means clustering analysis^39^ was performed to calculate the thickness of the hybrid dentin-resin layer (**Figure 4g** – dentin (orange), hybrid layer (grey), resin (blue)). Resin materials with BMEP show a hybrid layer thickness of 22 *μm*, compared to 20 *μm* for samples without BMEP (**Figure 4h**). This hybrid layer thickness is larger than the numbers reported for other resin systems, which is from 0.3~10 *μm*^40–43^. These results suggest the addition of BMEP increases the thickness of the hybrid layer by 10%, which improves the mechanical stability of the dentin-resin interface. This is consistent with the shear strength results and other studies that suggest the importance of reinforcing the hybrid layer ^40, 44^. This nanoindentation result shows consistency in a more stabilized hybrid layer between materials to enhance bonding strength^41^.

Biocompatibility tests were performed using resin-treated conditioned media to confirm the triacrylate-trithiol polymer adhesive system supports cell viability^10^ (**Figure 5 & S6**). Fully cured resin samples with and without BMEP were conditioned in low glucose DMEM at 37°C. Media was collected after 7 days to treat in BJ fibroblast cells. Cell morphology was imaged after 24 h exposure to original condition media by fluorescence imaging of cells stained for cell nuclei (DAPI, blue) and F-actin (phalloidin, green) (**Figure 5a & S6**). The cytotoxicity of resin materials was measured by LDH release after 24 h exposure to serially diluted original condition media. In the 100% condition media group, the relative cell viability of resin with BMEP was 83.24%, and the resin without BMEP was 80.97%; in the 50% and 25% condition media group, the relative cell viability of resin with BMEP was 86.33% and 86.39%, respectively, compared to 88.54% and 87.71% for the resin without BMEP (**Figure 5b**). These revealed low cytotoxicity of this triacrylate-trithiol polymer system and no significant differences between the resins with and without BMEP. BJ cells were retrieved and cell numbers in each condition were counted after 24 h exposure to the serially diluted condition media. Media containing released residuals of resin materials with and without BMEP did not significantly affect cell number and proliferation (**Figure 5c**). Together, we confirmed that TMPTA resin systems had low cytotoxicity, and resins with and without BMEP didn’t exhibit significant differences in their biocompatibility. This finding can be applied to incorporate additional functional groups for drug delivery and remineralization at the mineralized tissue interface.

**Figure 5.**
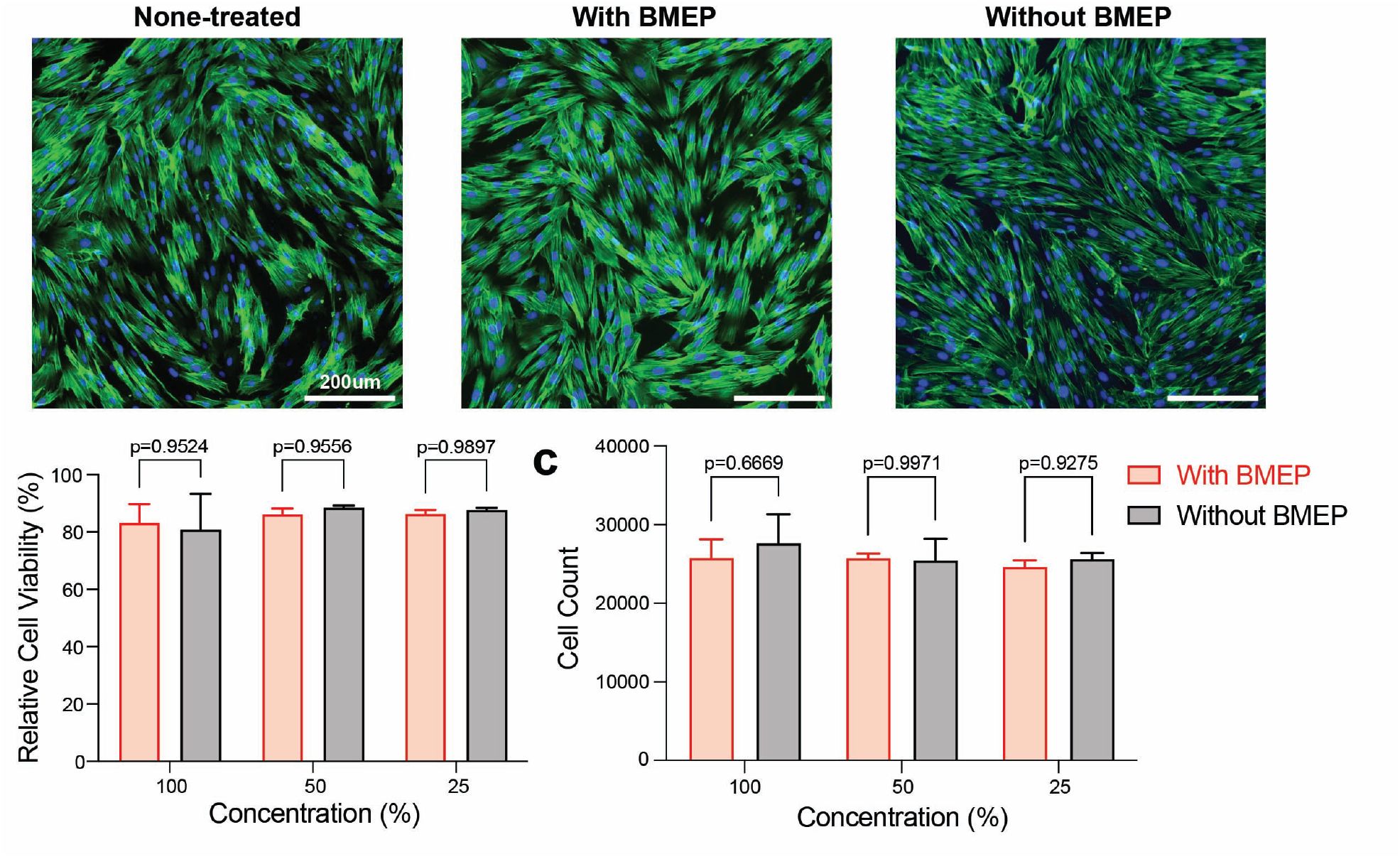
Biocompatibility of resin systems with and without BMEP. **(a)** Fluorescence imaging (scale bar 200 *μ*m) of BJ cells after 24 h culture in original condition (100% concentration) media compared to negative control, stained for nuclei (blue) and F-actin (green). **(b)** Relative cell viability, compared to negative control, and **(c)** cell counts of BJ cells after 24 h culture in condition media. Condition media are diluted with DMEM to a total concentration of 100%, 50% and 25% of the original condition media. n = 3 biological replicates, error bars represent SD.

## Conclusion

This study demonstrates the mechanical and adhesive properties of a multi-functional polymeric resin system composed of TMPTA and BMEP, designed to improve adhesion to mineralized tissues. Molecular dynamics simulations identified 2.5:1 TMPTA: TMPMP as the optimal ratio for the thiol-ene polymerization reaction, which was experimentally validated through rheological and mechanical testing. The addition of BMEP in the TMPTA resin played a crucial role in enhancing adhesion strength, achieving a shear bond strength of 10.8 MPa, comparable to commercial primer Clearfil SE Bond, without significantly affecting biocompatibility or shear modulus. Nanoindentation mapping further revealed that BMEP increased dentin-resin hybrid layer thickness with a sigmoidal modulus profile. Biocompatibility tests demonstrated that the TMPTA resin materials with and without BMEP had low cytotoxicity. The combination of micromechanical interlocking and chemophysical bonding with a thiol-ene polymer network enables strong adhesion while maintaining biocompatibility. Overall, the TMPTA-TMPMP-BMEP resin system presents a promising strategy for improving the longevity and clinical performance of dental adhesives, with potential applications beyond dentistry, including bone fixation and biomedical coatings.

## Data Availability

All data to generate figures for this manuscript will be made available prior to publication in a public repository.

## Author contributions

Authors YL, KTT, and KHV contributed to the conceptualization, data curation, writing, and review & editing of this manuscript. YL, CZ, SF, JL, KC, YJ and OF contributed to the data collection and analysis. YL, RX, and OAT contributed to the protocol formation. YH and LR contributed to the data modeling and visualization. All authors contributed to review and editing.

## Acknowledgments

This work was supported by the Joseph and Josephine Rabinowitz Award for Excellence in Research from Penn Dental Medicine. KHV received support from the National Institutes of Health for this work, NIH/NIDCR R00DE030084. This work was carried out in part at the Singh Center for Nanotechnology, which is supported by the NSF National Nanotechnology Coordinated Infrastructure Program under grant NNCI-2025608. We would like to acknowledge the undergrads from University of Pennsylvania, Hyunil Kim, Asset Yermekkaliyev, Sylvia Chen, Justin Wang and their advisor Shu Yang, who helped on the initial material synthesis protocol. We thank Luc Capaldi and Kailin Chen for their invaluable training and support on Nanoindentation test. And we also thank Brandley Tanelus for the preparation of dentin sample for shear bond test. Their assistance was instrumental in the successful completion of this research. LR acknowledges that this work was conducted during her sabbatical leave from CMU at the University of Pennsylvania.

