## Supplementary Information for "Biocompatible Multi-functional Polymeric Material for Mineralized Tissue Adhesion"

### **Supplemental Information**

### **Nanoindentation Test**

The resin film was crafted by pipetting 100μL of resin solution onto one MTC Bio microscope slide with a 25*75mm dimension. This initial layer was then complemented by carefully stacking another slide in a cross formation, creating a square area (25*25mm) with the resin solution sandwiched between them. Next, a UV light with an intensity of 103.2mW/cm² was directed onto the square area for a duration of 60 seconds using a UV lamp. Once the resin film had undergone complete curing, it was gently peeled off from the slides, designating the side where the light entered as the front side, and the side where the light exited as the back side. Subsequently, seven different points, spaced at 8 mm intervals, were chosen on both sides for the nanoindentation test.

Nanoindentation experiments were carried out using KLA iMicro Nano indenter with a Berkovich tip and the advanced E&H method. This methodology involved setting specific parameters: a target load of 10 mN, a target depth of 1000 nm, and a target indentation strain rate of 0.1/s. All tests were carried out at room temperature (22–24 °C).

### **Supplemental Figures**

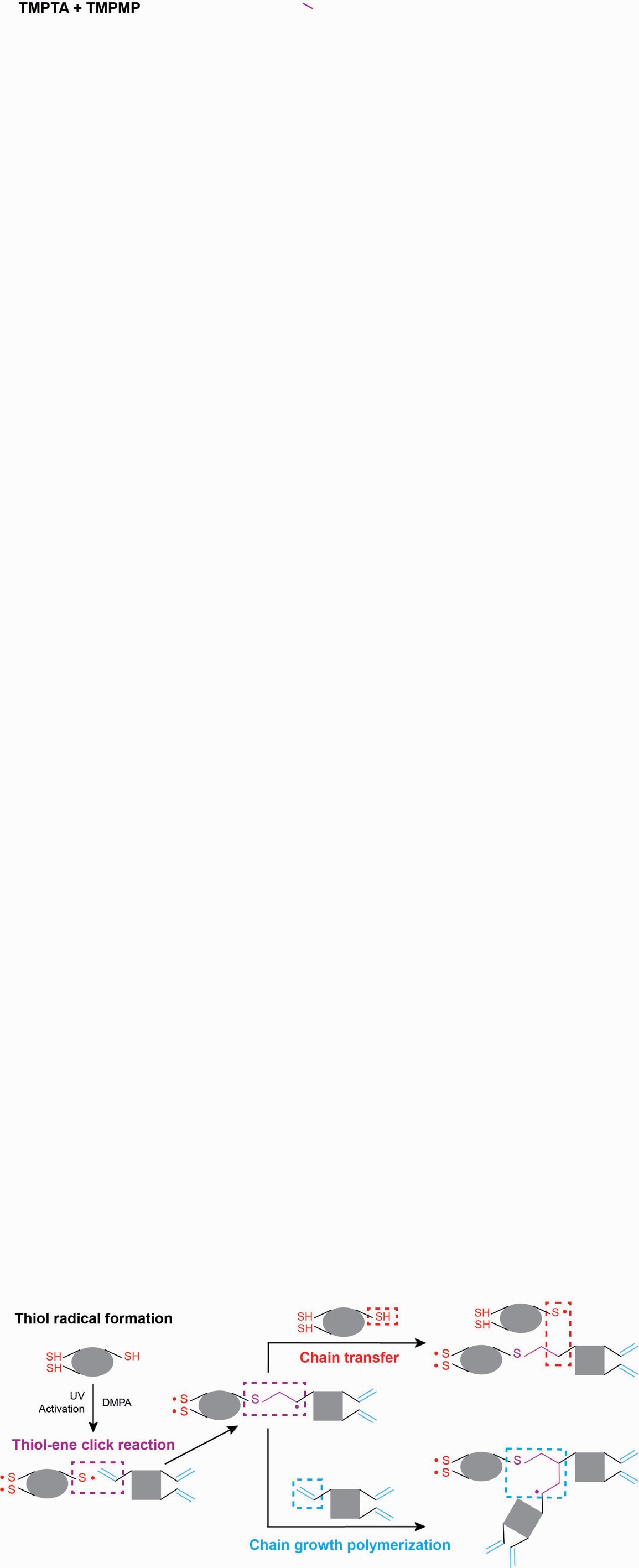

Figure S1. Thiol-ene click reaction processes for crosslinking TMPTA and TMPMP.

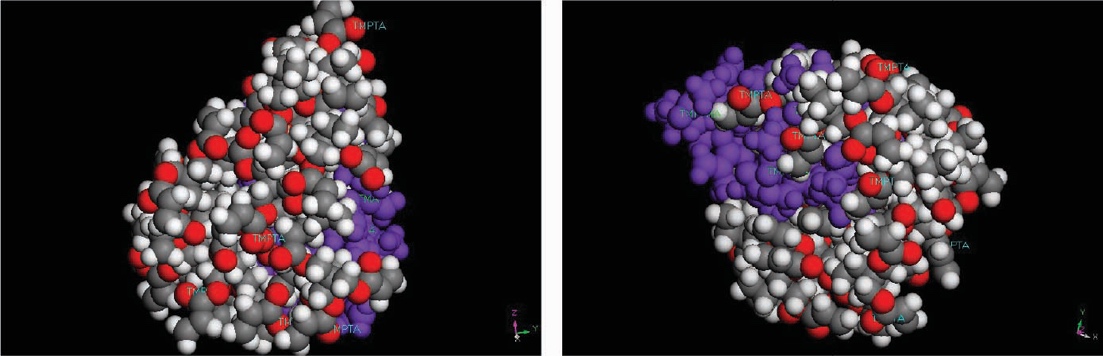

Figure S2. The predicted molecular structure of the dental resin material with the five TMPTA molecules and two TMPMP molecules. TMPTA: red – oxygen atom, grey – carbon atom, white – hydrogen atom; purple – TMPMP molecule.

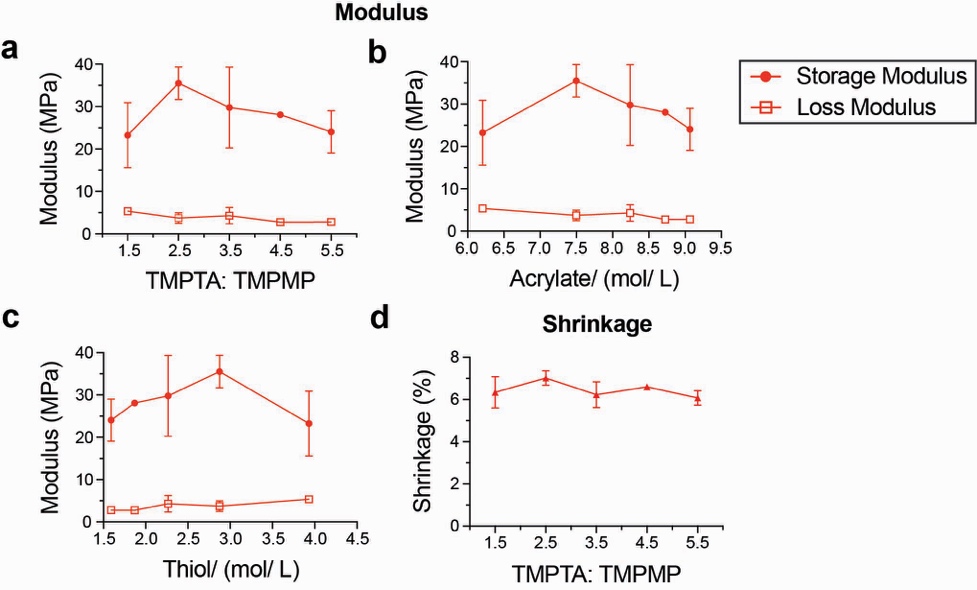

Figure S3. Storage modulus and loss modulus of resin with BMEP plotted as a function of: (a) TMPTA: TMPMP ratios, (b) acrylate concentration and (c) thiol concentration. (d) Shrinkage of resin with BMEP (5 wt%) under different TPMTA: TMPMP ratios.

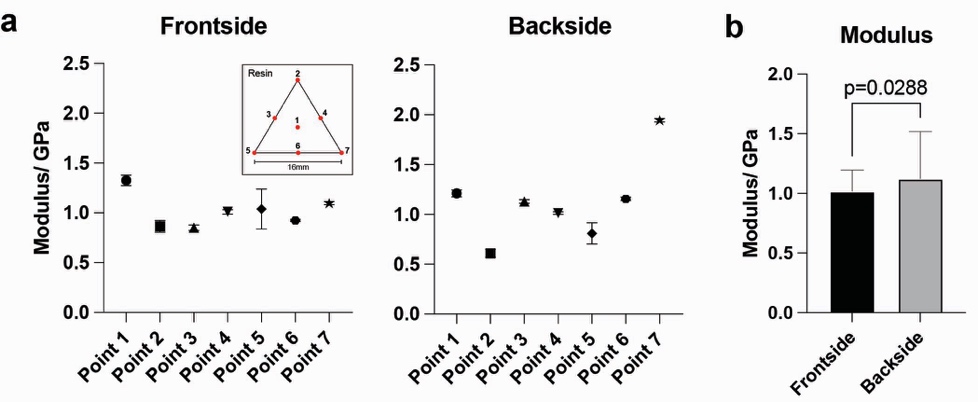

Figure S4. (a)Modulus data from Nanoindentation on thin resin film’s frontside and backside. (b)Statistics analysis of t test on modulus data of frontside and backside (sample size=77).

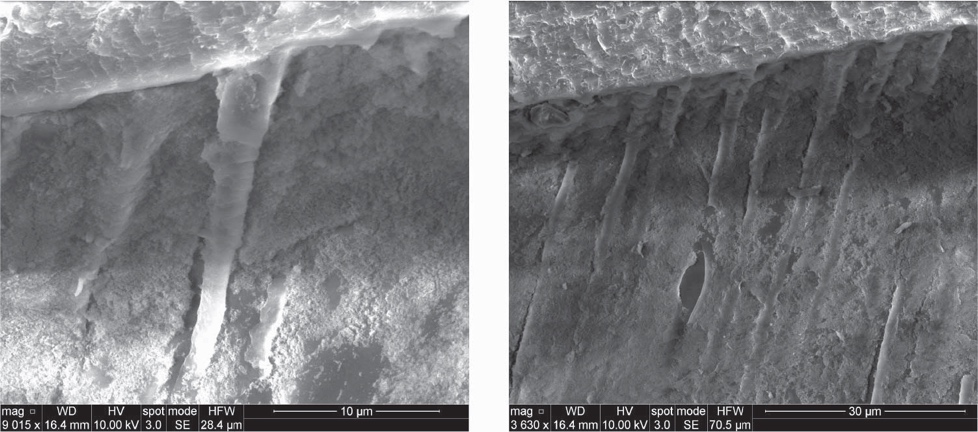

Figure S5. SEM images of along-tubule-resin-coated dentin interface with resin tag exposed under 10 $\mu$m and 30 $\mu$m scale.

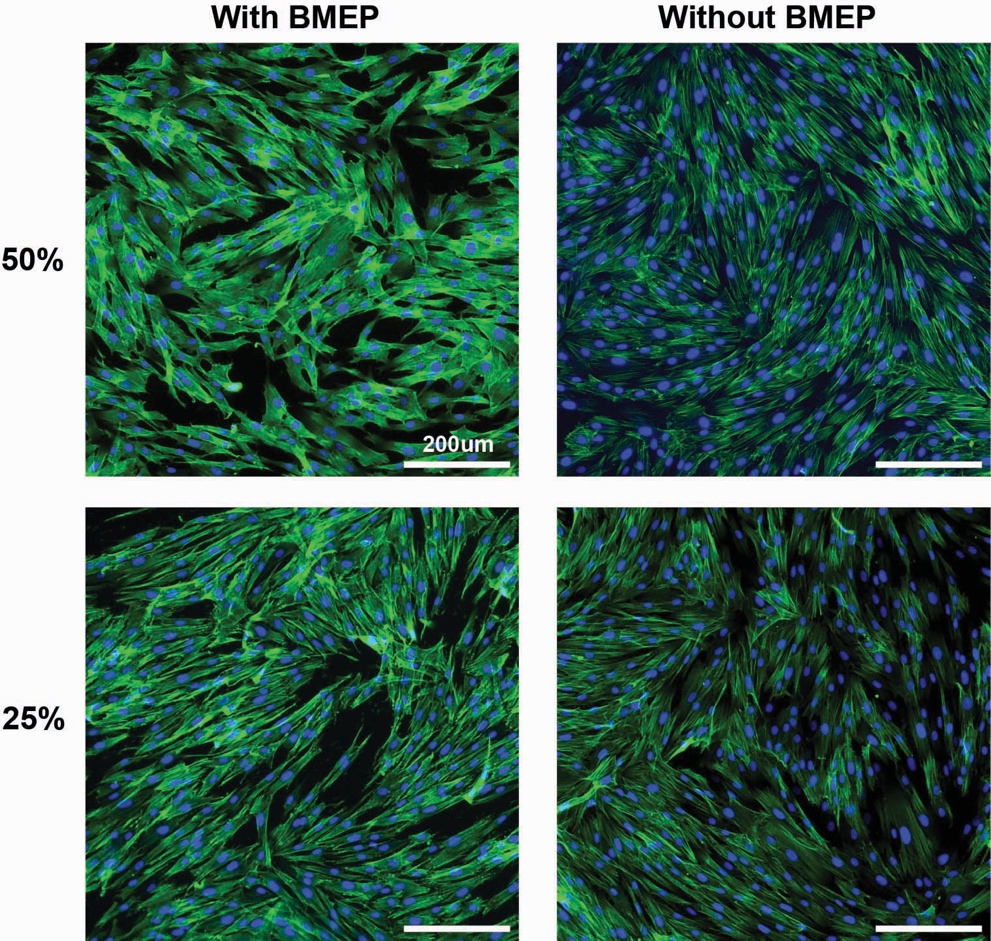

Figure S6. Fluorescence imaging of BJ fibroblasts treated 24 hours with conditioned media (25 and 50% dilutions) from resin samples with or without BMEP. F-actin (green, phalloidin) and nuclei (blue, DAPI). Scale bar 200 micrometers.
